# SARS-CoV-2 proteins bind heme and hemoglobin

**DOI:** 10.1101/2021.04.16.440124

**Authors:** Guilherme Curty Lechuga, Franklin Souza-Silva, Carolina de Queiroz Sacramento, Monique Ramos de Oliveira Trugilho, Richard Hemmi Valente, Paloma Napoleão-Pêgo, Suelen da Silva Gomes Dias, Natalia Fintelman-Rodrigues, Jairo Ramos Temerozzo, Nicolas Carels, Carlos Roberto Alves, Mirian Claudia de Souza Pereira, David William Provance, Thiago Moreno Lopez Souza, Salvatore Giovanni De-Simone

## Abstract

The coronavirus disease 2019 (COVID-19) pandemic, caused by severe acute respiratory syndrome virus 2 (SARS-CoV-2), has led to a global crisis that included collapsing healthcare systems and shut-down communities, producing considerable economic burden. Despite the number of effective vaccines quickly implemented, the emergence of new variants is a primary concern. The scientific community undertook a rapid response to better study this new virus. However, critical questions about viral protein-protein interactions and mechanisms of its physiopathology are still unclear. Although severe COVID-19 was associated with hematological dysfunctions, scarce experimental data were produced about iron dysmetabolism and the viral proteins’ possible interaction with hemoglobin (Hb) chains. This work demonstrates the binding of SARS-CoV-2 proteins to hemin and Hb using a multimethodological approach. *In silico* analysis indicated binding motifs between a cavity in the viral nucleoprotein and hemoglobin’s porphyrin coordination region. Different hemin binding capacities of mock and SARS-CoV-2-infected culture extracts were noticed using gel electrophoresis and TMB staining. Hemin-binding proteins were isolated from SARS-CoV-2-infected cells by affinity chromatography and identified by shotgun proteomics, indicating that structural (nucleoprotein, spike, and membrane protein) and non-structural (Nsp3 and Nsp7) viral proteins interact with hemin. *In vitro* analyses of virus adsorption to host cells and viral replication studies in Vero cells demonstrated inhibitory activities - at different levels - by hemin, protoporphyrin IX (PpIX) Hb. Strikingly, free Hb at 1μM suppressed viral replication (99 %), and its interaction with SARS-CoV-2 was localized to the RBD region of the Spike protein. The findings showed clear evidence of new avenues to disrupt viral replication and understand virus physiopathology that warrants further investigation.

## Introduction

At the end of 2019, severe acute respiratory syndrome coronavirus 2 (SARS-CoV-2) was first recognized in Wuhan (Hubei province, China). The disease rapidly spread to many countries due to its high transmissibility and prolonged incubation, allied to the existing highly connected global travel network [1]. This zoonotic virus became the etiological agent of the 2019 coronavirus disease (COVID-19). As an ongoing pandemic disease, COVID-19 has proven to be a significant economic and public health challenge. The elderly and individuals with pre-existing comorbidities are severely affected, but severe COVID-19 can impact the full range of age groups [2]. The global scientific community has exerted tremendous efforts to understand the viral structure and physiopathology to identify control measures that include drug repurposing strategies, plasmapheresis, and vaccination [3]. Despite the performance and increasing availability of the newly developed vaccines, the recent detection of emerging variants that appear to escape from the immune responses represents a significant concern to immunization strategies [4].

Although drug repurposing has not been proved to be unequivocally satisfactory against SARS-CoV-2, this strategy could still be worth fighting COVID-19 when appropriate biochemical interactions of viral proteins and small molecules are determined. Possibly, biochemically-based evidence of effective treatments for current and future variants can be accelerated by expanding the breadth of knowledge on the activities of the individual SARS-CoV-2 proteins during infection, avoiding further frustration in clinical trials with molecules with limited preclinical effectiveness against SARS-CoV-2, such as lopinavir (targeting viral protease) in combination with ritonavir (LPV/RTV), and hydroxychloroquine (HCQ) [5]. Hematological COVID-19 is a constitutive component in critically ill patients [6,7]. The heme-iron dysregulation has been observed in COVID-197, with binding signatures including hyperferritinemia, low hemoglobin (Hb) levels, low serum iron, anisocytosis, and increased variation of red blood cell distribution width (RDW), and hypoxemia [8–10]. Unbalanced erythrocyte counts, Hb, and iron levels were associated with poor clinical outcomes in COVID-199. An *in silico* analysis pointed to a relevant role of SARS-CoV-2 proteins in viral physiopathology. The predictions suggest that the capture of heme, resulting from a coordinated attack of orf1ab, ORF10, and ORF3a to the 1-β chain of hemoglobin, could interfere with heme metabolism and oxygen transport. This analysis also proposed the binding of heme by structural and non-structural proteins of SARS-CoV-2 [6]. Although these data bring interesting perspectives, experimental confirmation is still needed.

Heme, iron protoporphyrin IX (PpIX), is a ubiquitous molecule with importance in numerous biological processes such as a cofactor for proteins (Hb), transcriptional regulation [11], RNA processing [12], oxidative stress [13], inflammation [14], and coagulation [15], which are all critical aspects of COVID-19 pathology. Heme and porphyrins can modulate viral infection by targeting both viral structures and cellular pathways. Porphyrins have broad activities against different viruses such as hepatitis B virus (HBV), hepatitis C virus (HCV), human immunodeficiency virus (HIV), and Zika virus (ZIKV) [16,17]. Nonspecific heme interactions, including hydrophobic binding to viral surface envelope proteins, block viral cellular entry [18]. Potent antiviral activity of PpIX and verteporfin in the nanomolar range has been recently reported in the inhibition of viral invasion by blocking the virus-cell fusion mediated by SARS-CoV-2 Spike (S) protein and ACE2 [19]. When Vero cells were pretreated with both, there was an inhibition of viral RNA production, suggesting that their interactions with ACE2 caused the viral entry block.

Despite the concerted efforts to unveil key viral targets, experimental evidence is still limited. Here, we focused on the capacity of SARS-CoV-2 proteins to capture heme and Hb. Our findings demonstrate that SARS-CoV-2 structural and non-structural proteins can bind to hemin. An *in silico* analysis identified heme-binding motifs in nucleoprotein and *in vitro* assays showed that the approved drugs hemin and PpIX precluded - at different levels - viral attachment to host cells, reducing viral replication. Notably, free Hb suppressed viral entry by interaction with the RBD region in the Spike protein and reduced viral replication. The results suggest that these molecules could be promising candidates for the treatment of COVID-19 and highlight the need to investigate further the mechanisms involved with the iron dysmetabolism observed in infected individuals.

## Results

### In silico analysis

The mapping of the binding motifs to heme was the first step in this study’s *silico* approach. Analysis of 68 crystallized hemoglobin/heme complex structures (**Supplementary Table 1**) identified conserved amino acid residues (identical in at least 97 % of the analyzed structures in the same position) that interact with the heme group. Furthermore, Histidine displayed the highest binding frequency to heme, while Alanine had the lowest contribution (**Fig. 1a**). From the mapped motifs in human hemoglobin (**Supplementary Figure 1**) three equivalent motifs in SARS-CoV-2 nucleoprotein - presenting different E-values (**Supplementary Figure 2**): motif 1 = 3.4 × 10^−505^, motif 2 = 4.4 × 10^−532^ and motif 3 = 1.8 × 10^−628^ - were identified (Fig. 1b). These analyses also indicate a correspondence regarding the residues Tyr (in motif 1), Lys (in motif 2), and Leu and Phe (in motif 3) in heme-binding motif composition between human Hb and the nucleoprotein of SARS-CoV-2 (**Fig. 1b**).

**Fig. 1:**
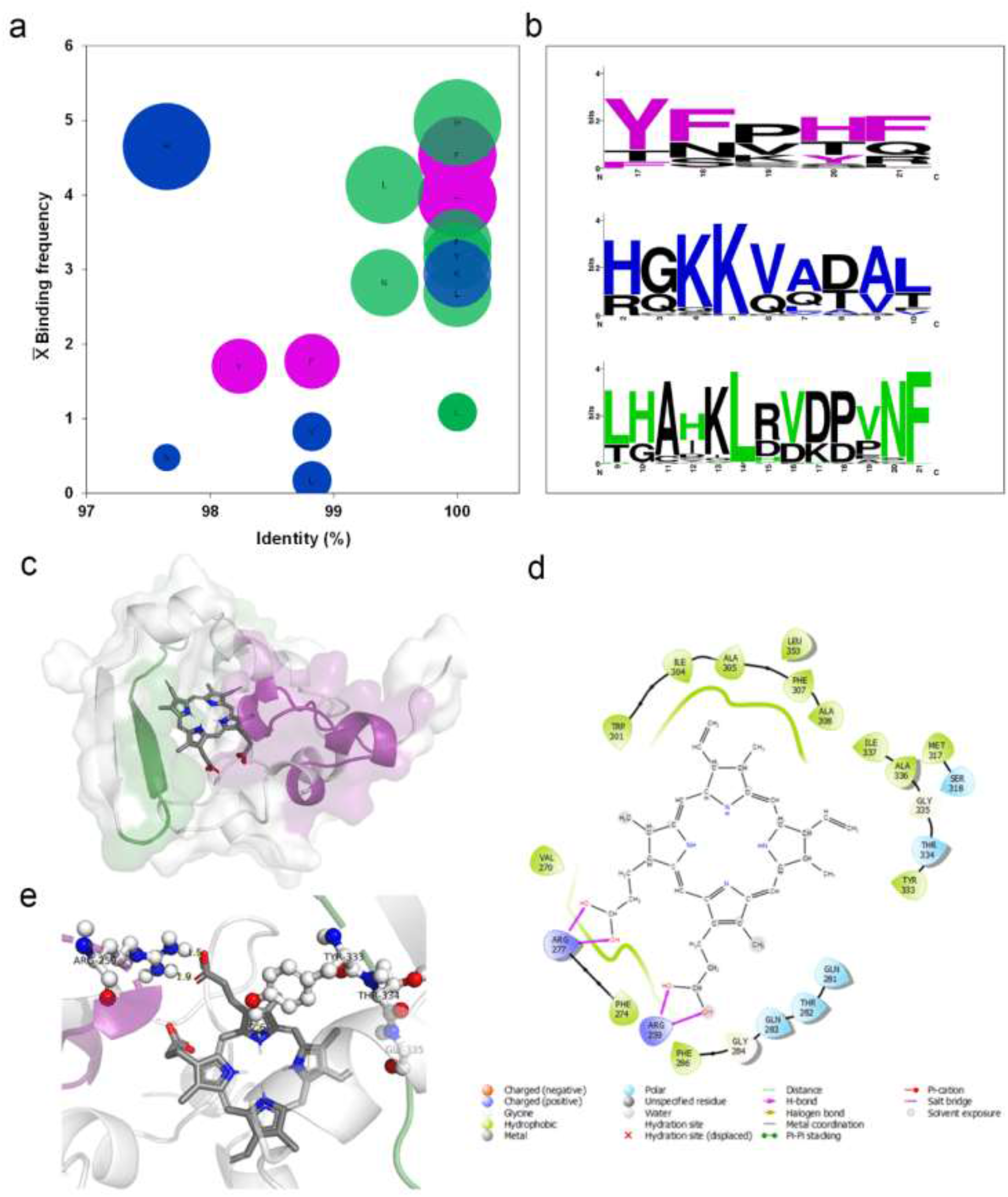
Heme-binding motifs identified by an *in silico* analysis. (**a**) the heme-binding motifs for 68 Hb structures deposited in the PDB data bank were mapped to define the frequency of amino acid occurrence (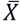 Binding frequency) and degree of identity (%). The balls’ size is relative to the number of connections (5 to 1), displayed from biggest to smallest in size. (**b**) three binding motifs were identified in the nucleoprotein of SARS-CoV-2 by an analysis in the MEME-Suite server of Hb amino acid sequence obtained from Uniprot sever. Colored amino acids related to residues in Hb that interact with heme. Black residues form part of the motif without binding heme. The size of the amino acid relates to the number of occurrences (bits). Molecular docking of SARS CoV-2 nucleoprotein with human PpIX. (**c**) Binding position of PpIX (sticks) indicating the orientation of binding with nucleoprotein (surface) to predicted motifs: purple = motif 1 and green = motif 3. (**d**) 2D representation indicating types of bonds that occur in nucleoprotein pocket bounded with PpIX. (**e**) 3D model showing nucleoprotein amino acid residues (ball and sticks) that compose motifs 1 and 3 coordinating the binding with the PpIX (bars).

**Fig. 2:**
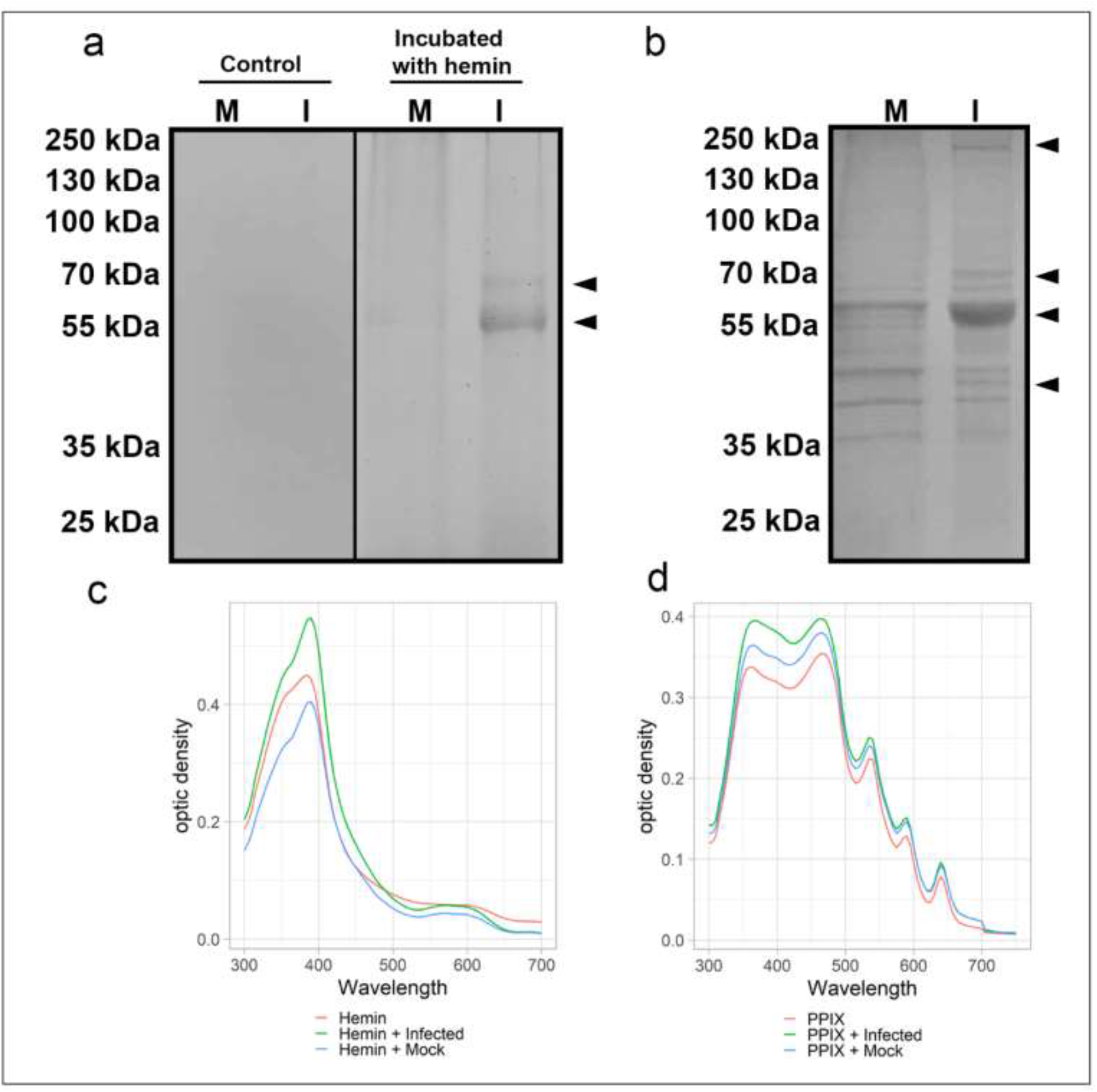
Heme-binding proteins in extracts from SARS-CoV-2 infected and non-infected Vero cells along with a spectroscopic binding analysis in solution. Total protein extracts (20 µg) from virus-infected (I) and mock-infected (M) were: **a**) incubated with 300 µM of hemin for 1 h at 25 °C, separated by SDS-PAGE, renatured and exposed to TMB in-gel to reveal hemin or (**b**) resolved by SDS-PAGE, transferred to nitrocellulose membrane and incubated with hemin (2 µM) for 1 h before revealing hemin-protein complexes by DAB. Arrowheads indicate bands of hemin-protein complexes. UV-visible spectra of hemin (**c**) alone (10 µM, red) or in total protein extracts (20 µg) of virus-infected Vero cells (green) and mock-infected cells (blue). UV-visible spectra of PpIX (**d**) alone (5 µM, red) or in total protein extracts (20 µg) of virus-infected Vero cells (green) and mock-infected cells (blue). Changes in the Soret peak and Q-bands are observable. Data are representative of two independent experiments. Source data are provided as a Source Data file.

The molecular docking assay was initially necessary to identify possible binding pockets in the SARS-CoV-2 nucleoprotein. A cavity with a -9.8 kcal/mol binding energy and a 121 Å size was selected to be assayed. In this cavity, it was possible to identify the bonds of PpIX with motifs 1 and 3 (**Fig. 1c**), with a predominance of hydrophobic bonds followed by polar bonds and hydrogen bonds (**Fig. 1d**) with porphyrin propionate groups. Both motifs’ potential to contribute to the coordination of protoporphyrin binding was related with Arg259 of motif 1 performing hydrogen bonding and Tyr333, Thr334, and Gly335 of motif 3 performing hydrophobic and polar bonds, respectively (**Fig. 1e**).

### Proteins from SARS-CoV-2-infected cells extract bind to hemin (heme)

Two distinct approaches evaluated the cellular and viral proteins’ interaction with heme. In the first, 300 µM hemin was pre-incubated with protein extracts obtained from Vero cell cultures, either infected with SARS-CoV-2 or mock-infected, as a control, followed by SDS-PAGE separation, in-gel protein renaturation, and heme-binding protein visualization using TMB reaction. Two protein bands (*ca*. 55 and 70 kDa) were revealed for SARS-CoV-2 infected Vero cells protein extract sample, with only a very faint band stained at ∼55 kDa in the mock protein extract sample **(Fig. 2a; right side**). As expected, no bands were visualized from the oxidation of TMB oxidation for the negative control samples in the absence of hemin’s addition (**Fig. 2a; left side**). The alternative approach separated protein extracts by SDS-PAGE under denaturing and reducing conditions transferred to a nitrocellulose membrane before incubation with 2 µM hemin. Bound hemin was revealed by its reaction with DAB. This approach displayed a greater sensitivity for detecting protein-heme complexes, as seen by the more significant number of visible protein bands. The patterns showed differences between the proteins’ profiles that can interact with heme in uninfected and virus-infected cells. The bands previously observed by the first approach were also detected, with the band at ∼55 kDa being the most prominent. Among the revealed bands, four (*ca*. 45, 55, 70, and 230 kDa) were exclusively detected in the protein extract of SARS-CoV-2-infected cells (**Fig. 2b**).

These observations of heme-binding protein in-gel and on membranes raised questions about their binding properties in solution. UV-visible spectra of hemin (10 µM) diluted in PBS were recorded to analyze possible viral protein interactions with heme, revealing a Soret peak with a lambda maximum at 385 nm. Changes in hemin absorption spectra were observed after incubation with mock and virus-infected protein extracts. A redshift was noticed in both cellular extracts (lambda max. 390 nm) (**Fig. 2c**). Furthermore, the Soret band and protoporphyrin IX absorbance in both the Soret region and the Q bands were increased in mock and virus-infected cell extract (**Fig. 2d**).

### SARS-CoV-2 proteins identification and confirmation of binding to hemin and hemoglobin

Shotgun proteomics was used to identify which SARS-CoV-2 proteins were interacting with heme. Initially, SARS-CoV-2-infected Vero cells total protein extract was subjected to SDS-PAGE separation, followed by excision of all visible bands and submission to processing for protein content identification nanoelectrospray coupled to high-resolution tandem mass spectrometry (LC-MS/MS). This approach yielded virus protein identifications in the six bands indicated in figure 3a. Nucleoprotein was identified in almost all excised bands (2, 3, 4, 5, and 6). Spike protein was identified in bands 1, 2, 3, and 6, while membrane protein was identified in bands 1 and 3. The non-structural proteins NSP3 and NSP2 were detected in bands 1 and 3, respectively (**Fig. 3a**). The most abundant protein identified in band 4 was albumin, probably due to supplementation in the culture medium. To reduce the amount of this contaminant protein that could interfere or mask the signal of viral proteins, albumin was depleted using affinity chromatography followed by hemin-agarose binding. Next, albumin-depleted protein extract of SARS-CoV-2 infected Vero cells was incubated with hemin-agarose beads to confirm that viral proteins can interact and bind heme. Finally, the hemin-agarose eluate containing the hemin-binding proteins was subjected to SDS-PAGE under denaturing and reducing conditions, revealing the presence of several viral proteins, as identified by LC-MS/MS (**Fig. 3b**). In-band 7, nucleoprotein, Spike, Nsp3, Nsp7, and membrane protein were placed. Nucleoprotein was also present in band 8 with the highest spectral counts and Spike protein in lower abundance. Additional information regarding cellular proteins and spectral counts obtained for each band can be found in the supplemental material (**Supplementary Data 1**). Additionally, after the interaction, western blot - with immunostaining using convalescent patient serum - revealed two reactive bands at ∼70 kDa and ∼55 kDa in the total extract, one at ∼70 kDa in the unbound fraction, and two bands (one intensely reactive at ∼55 kDa and the other at ∼45 kDa) in the bound fraction (**Fig. 3c**).

**Fig. 3:**
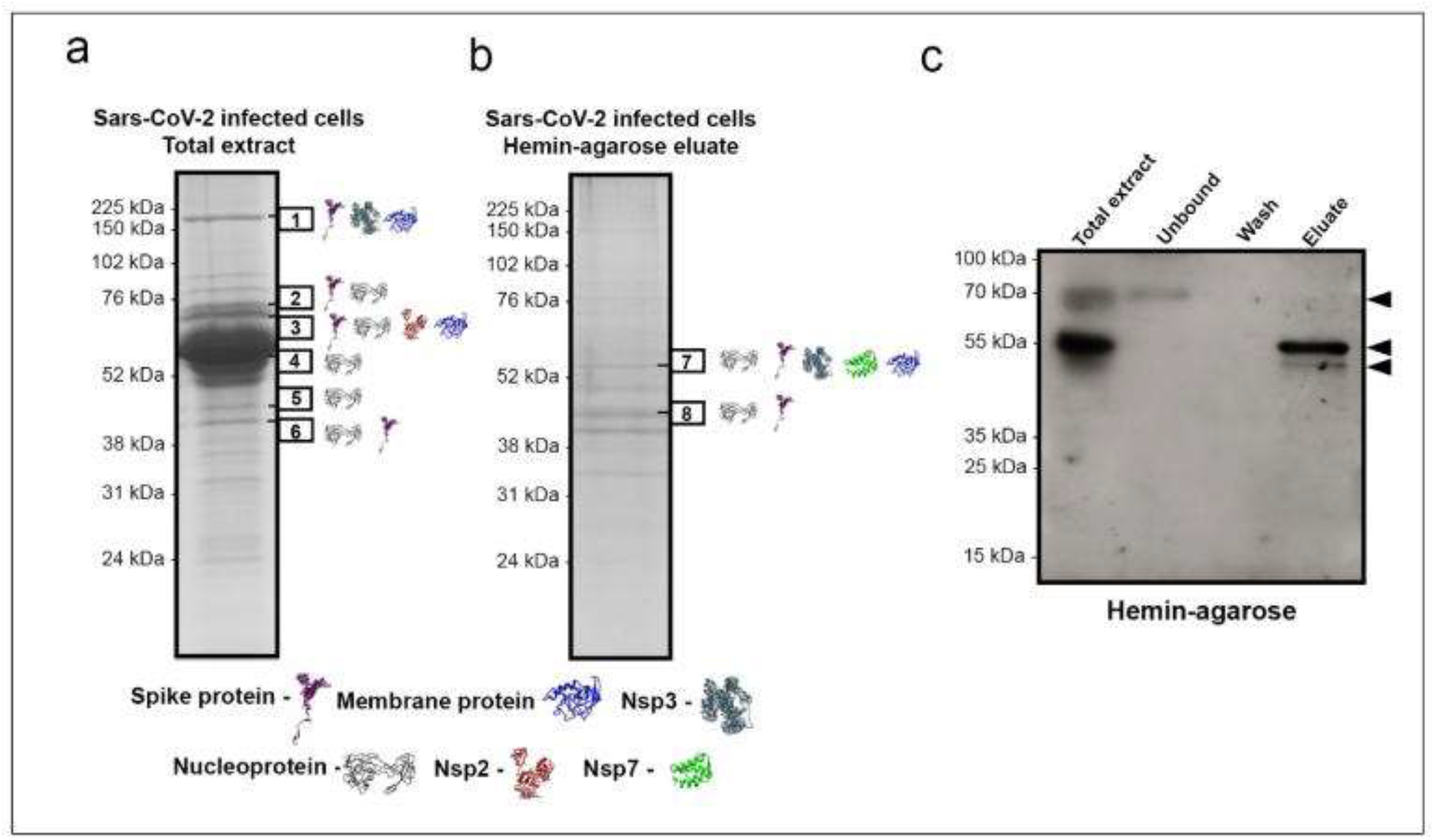
Heme affinity purification of proteins from total extracts of SARS-CoV-2 infected Vero cells and their identification by mass spectrometry. SDS-PAGE separated proteins stained by Coomassie blue from (a) whole protein extract of virus-infected Vero cells and (**b**) eluate from a hemin-agarose purification. Bands excised for protein identification by LC-MS/MS are indicated by black boxes 1-8. The proteins identified were nucleoprotein (P0DTC9), membrane protein (P0DTC5), Spike protein (P0DTC2), and Replicase polyprotein 1a/1ab (P0DTC1/P0DTD1). The peptides in the replicase polyprotein 1a/1ab were identified as Nsp2, Nsp3, and Nsp7. The identified proteins in each band are shown as 3D structures retrieved from the I-Tasser site (https://zhanglab.ccmb.med.umich.edu/I-TASSER/). (**c**) Western blot with immunostaining - using COVID-19 patient convalescence serum to detect reactivity - from hemin-agarose chromatography fractions of protein lysate from SARS-CoV-2-infected Vero cells; arrowheads indicate two bands revealed in the eluate.

The observation that viral proteins can bind to hemin suggested that this interaction could extend to Hb. To test this hypothesis, an overlay assay was performed with Hb on SARS-CoV-2 infected Vero cell extracts separated by SDS-PAGE and transferred to a nitrocellulose membrane (**Fig. 4a)**. Haptoglobin, used as a positive control, displayed the predicted size for the β-chain at 40 kDa and showed that this approach could detect Hb binding proteins; it revealed two protein bands (∼150 kDa and ∼75 kDa) exclusively in SARS-CoV-2-infected cells, while the other bands (∼95 kDa, ∼70 kDa, and ∼50 kDa) were also found in the mock cell extract. Based on the results from mass spectrometry, the bands’ sizes suggested the Spike protein presence (Fig. 4a). To confirm this conclusion and refine the region in Spike glycoprotein that could interact with Hb, a protein-protein interaction assay was performed using the Spot Synthesis technique. An array of 15-mer peptides with a 5 amino acid overlap representing the RBD region of the Spike protein synthesized *in situ* on a cellulose membrane was constructed. Following Hb’s incubation and its subsequent detection by Hb-specific antibodies with corresponding secondary antibodies, several highly reactive spots were revealed that indicated the natural motifs interaction between sequences in the RBD and Hb (**Fig. 4b, top panel**). The relative intensity percentage was calculated, and individual peptide sequences were identified; signal intensities above 50% were considered the cutoff for a positive reaction. Five unique peptide sequences were defined that interacted with Hb (**Fig. 4b, lower panel**). Molecular docking assay of protein/protein interactions indicated the possibility of binding the spike protein with the alpha and beta hemoglobin domains (**Fig. 4c**), showing binding energy of -460 kcal/mol (**Supplementary Figure 3**). The data showed that this binding could be coordinated by four hot spot amino acid residues (**Supplementary Table 2**) from both proteins: Spike (Tyr341, Tyr351, Phe347, Arg346, Tyr451, and Leu452) and Hb (His46, His51, Gln55, and Asp48) (**Fig. 4d**).

**Fig. 4:**
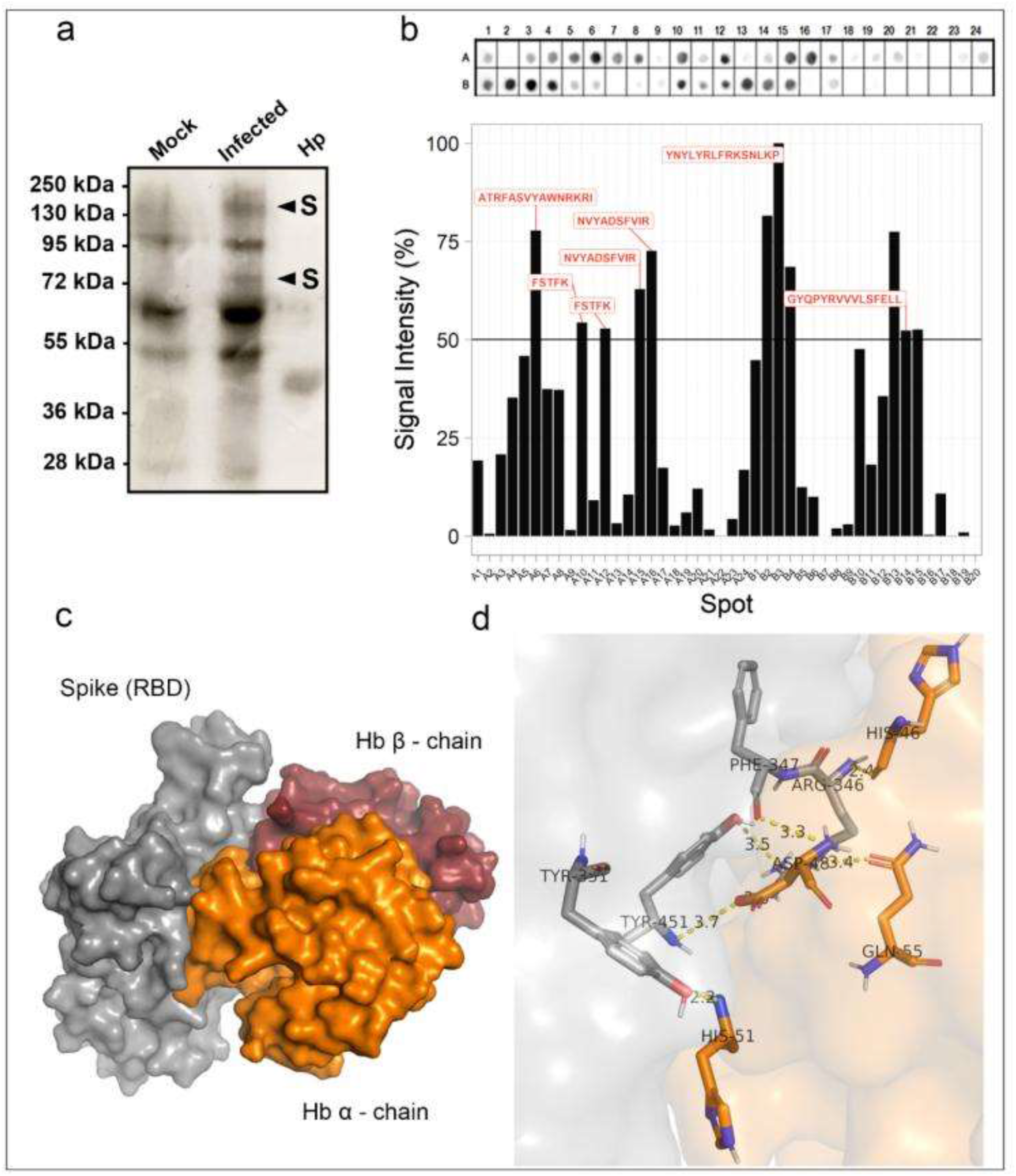
Hemoglobin (Hb) binding to Spike protein and its RBD. (**a**) Total protein extracts (20 µg) of SARS-CoV-2-infected Vero E6 cells (Infected) or mock-infected cells (Mock; 20 µg) were separated by SDS-PAGE (10%) along with haptoglobin (Hp; 2 µg), transferred to a nitrocellulose membrane, and serially incubated with Hb (10 µg/mL), anti-Hb antibody, and peroxidase-conjugated secondary antibodies. After substrate, two 72 and 150 kDa bands were exclusively found in SARS-CoV-2 infected extract (arrowheads). (**b**) Spot synthesis analysis with a library of 15-mer peptides offset by five amino acids to represent the RBD region of the Spike protein synthesized directly onto a cellulose membrane followed by probing with Hb (5 μg/mL) and revealed by anti-human Hb antibodies. The top panel shows the chemiluminescent image of signals from peptides bound to Hb. The bottom panel shows a graph of the signal intensities normalized to the maximum signal. An intensity level above 50% defined Hb-reactive peptides. Molecular docking of SARS CoV-2 spike protein with human hemoglobin (n=1). (**c**) Interaction of spike protein (gray) with α-chain Hb (orange) and β-chain (red). (**d**) Representation of amino acid residues binding to spike protein (sticks and gray) with α-chain of hemoglobin (orange). Source data are provided as a Source Data file.

### Hemin, hemoglobin, and PpIX affect SARS-CoV-2 adsorption and replication

The intense interaction between peptides in the RBD of the Spike and localization of Hb in the molecular model suggested that the presence of free Hb could competitively interfere with the binding of SARS-CoV-2 to ACE2, its receptor on host cells. As such, we hypothesized that the addition of Hb could impair virus replication in an *in vitro* assay. To test this hypothesis, Vero cells were pretreated with a sub-optimal Hb concentration (1 µM) for 1 h at 37 °C before their exposure to the virus and then infected SARS-CoV-2 at MOI of 0.01. Alternatively, the virus was preincubated with 1 µM of Hb for 1 h before its addition to Vero cells for another hour at 37 °C. After 24 h, culture supernatants were collected, and the production of infectious virus particles was quantified by plaque assay. The pretreatment of Vero cells with Hb did not affect virus production (**Fig. 5a, left column**); however, the virus’s preincubation with Hb reduced approximately 52.0 ± 1.3 % of virus replication. Convalescent plasma was used as a positive control for viral neutralization and inhibited viral replication (100 %)(**Fig. 5b, left column**). Hemin and PpIX were also tested to evaluate if porphyrin-protein interactions could affect virus replication. Hemin displayed no or little effect in virus replication when used as a pretreatment of Vero cells or SARS-CoV-2, respectively (**Fig. 5a and 5b, center columns**). PpIX inhibited SARS-CoV-2 replication by 90.0 ± 1.8 and 100 % when used as a pretreatment of the cells or virus, respectively (**Fig. 5a and b, right columns**).

**Fig. 5:**
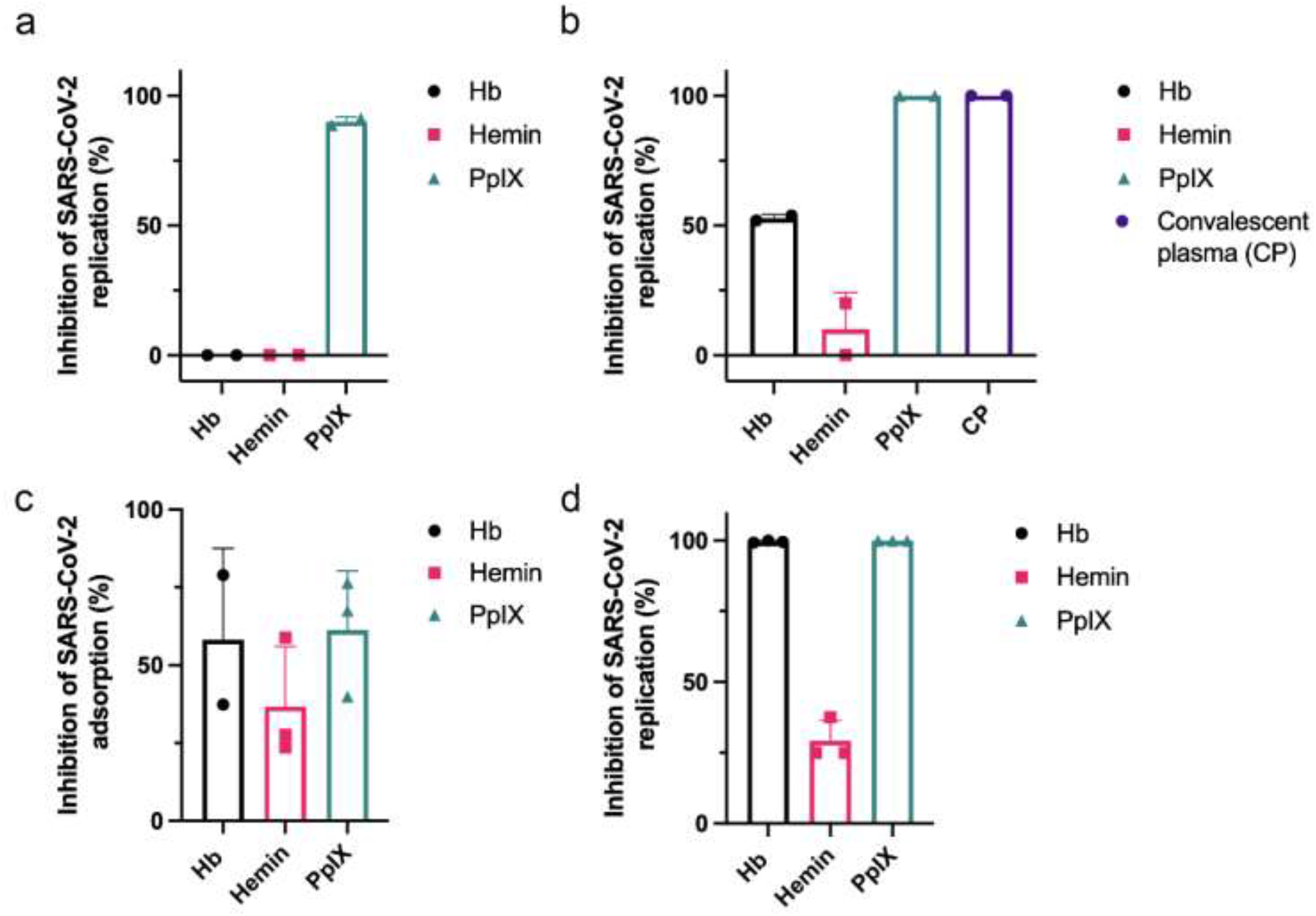
Inhibition of SARS-CoV-2 replication and attachment by porphyrins and Hb. (**a-b**) Vero E6 cells (**a**) or SARS-CoV-2 (**b**) were pre-incubated with 1 µM of Hb, hemin, or PpIX for 1 h at 37 °C before the start of infection with an MOI of 0.01 for an additional hour at 37 °C. Culture supernatants were collected after 24 h, and the virus titer was determined by plaque assay (n=2). A convalescent plasma of an infected patient (1:3 dilution) was used as the positive control. (**c**) SARS-CoV-2 was incubated at 1 µM Hb, hemin, or PpIX for 1 h at 37 °C before introducing Vero E6 cells at an MOI of 0.01 for 1 h at 4 °C. Rinsed cell monolayers were lysed, and RT-PCR quantified the virus content. Cumulative data (Hb: n=2, Hemin: n=3, PpIX: n=3). (**d**) Vero E6 was infected with SARS-CoV-2 at an MOI of 0.01 for 1 h at 37 °C and then treated with 1 mM of hemin, PpIX, or Hb. After 24 h, supernatants were collected, and the virus titer was quantified by PFU/mL (Hb, Hemin, PpIX: n=3). Data represented the percentage of inhibition compared to control (infected and untreated) and expressed as mean with standard deviation. Source data are provided as a Source Data file.

Next, we analyzed if viral replication reduction was related to Hb binding interference with a receptor-mediated virus attachment. An adsorption inhibition assay was performed. The incubation of the virus with cells was performed at 4 °C for 1 h to minimize internalization, and then viral RNA was purified for quantification by RT-PCR. When SARS-CoV-2 was pre-incubated with Hb (1 µM) for 1 h at 37 °C and then applied to Vero cells at an MOI of 0.01, a reduction of 58.1 ± 29.6 % was measured in the virus adsorption (**Fig. 5c**). Hemin (1 µM) reduced virus attachment by approximately 40 %, and PpIX reduced virus attachment by 61.3 ± 18.0 %. When cells were treated with 1 µM of either Hb, hemin, or PpIX, after the initiation of infection by SARS-CoV-2, virus replication was inhibited 99.5 ± 0.4 % by Hb, 100 % by PpIX, and 29.1 ± 7.2 % by hemin **(Fig. 5d**).

## Discussion

Severe COVID-19, caused by SARS-CoV-2, typically leads to pneumonia and acute respiratory distress syndrome (ARDS). However, the growing list of evidence indicates a systemic impairment that leads to multiorgan failure. During an infection, an imbalance in the immunological response can produce a “cytokine storm” and numerous other pathophysiological processes such as hypoxemia, thrombosis, pulmonary embolism, encephalopathy, myocardial injury, heart failure, and acute kidney injury20. Hematological dysfunction in severe COVID-19 includes low levels of erythrocytes and an increased variation in the red blood cell distribution width (RDW)8,9. Recent reports on the immune effects of COVID-19 have highlighted immune thrombocytopenia and autoimmune hemolytic anemia21. Other evidence suggests an increase in hemophagocytosis that could be related to elevated levels of ferritin in COVID-1922. In hemolytic disorders, the release of high amounts of Hb and heme trigger proinflammatory response, complement activation, and procoagulant and pro-oxidative environment [23,24].

Recently, an *in silico* analysis hypothesized that viral proteins could bind Hb beta-chains that would be expected to interfere with O2 transport and heme metabolism6. Another finding suggested that the sequence similarity between SARS-CoV-2 Spike protein and hepcidin, a peptide hormone involved in iron metabolism, could lead to an imbalance of iron metabolism7. Yet, in the absence of experimental data, it is not clear if SARS-CoV-2 proteins actually can interact with Hb and displace iron from heme.

To decipher these possible interactions, *in silico* analysis was performed to identify heme-binding motifs in viral proteins. Commonly, heme coordination can occur by hydrogen bonds via propionate groups, π-π stacking, electrostatic and hydrophobic interactions. The critical heme-coordinating amino acids are histidine, cysteine, and tyrosine; methionine and lysine occur at a lower frequency [25,26]. Here, *in silico* analysis supported the potential interaction of porphyrin ring with Tyr333 of the N protein in addition to hydrophobic and hydrogen bonds via propionate side chains. An open question is whether heme is a necessary endogenous cofactor recruited by viral proteins during replication and translation. The primary function of the N protein is to bind RNA. This protein comprises an N-terminal domain containing the RNA-binding site, a C-terminal dimerization domain, and a central linker region rich in serine and arginine [27]. The heme-binding motifs identified in the N protein suggest another function for this protein. The analyses of other N protein motifs identified a nuclear localization signal and a nuclear export signal that offer dynamic nuclear-cytoplasmic trafficking that can be involved in transcription regulation [27].

N protein binding to heme might have an important implication in viral and cellular transcriptional regulation since heme regulates multiple transcription factors in the nucleus, modulating the expression of various genes [28]. Interestingly, high-confidence protein-protein interactions between SARS-CoV-2 and human proteins identified N protein associated mainly with components of cellular translational machinery [29]. N protein was reported to interact with two subunits of casein kinase 2 (CK2), a protein involved in activation of heme regulator inhibitor (HRI) kinase that phosphorylates eukaryotic initiation factor 2 (eIF-2), inhibiting translation [30]. N protein could be involved in dissociation or recycling of heme from HRI, modulating the phosphorylation of eIF2-α [31]. Some viruses do not require eIF-2 and even induce host translation shutdown [32]; possibly, these events consist of a manipulation route for viral protein synthesis.

Electrophoresis of mock and SARS-CoV-2 infected cells revealed different heme-binding profiles. The electrophoretic mobility in the TMB in-gel staining assay suggests that Spike fragment (S1 and S2, ∼70kDa) and nucleoprotein (55 kDa) could bind hemin and retained its peroxidase activity. Also, heme-binding interactions in solution were changed by viral infection. Due to their hydrophobicity, heme and PpIX have low solubility in an aqueous solution and tend to aggregate. Binding complex with proteins can cause redshift and increase the Soret band’s intensity, and change spectra profile of Q band [33]. However, these results could also indicate a viral modulation of cellular hemeproteins or heme-binding proteins.

To refine the analysis, shotgun proteomics was performed to identify SARS-CoV-2 heme-binding proteins. Identification of proteins in SARS-CoV-2-infected Vero cells total extract revealed structural and non-structural proteins. Spike holoprotein was placed in the expected gel migration range (Band 1), but also cleavage products S1 and S2 produced by proteases (TMPRSS2, furin, cathepsin) were observed at ∼70 kDa bands (Band 2 and 3). Interestingly, SARS-CoV’s mass spectrometry analysis also identified Spike protein in bands of relative molecular masses divergent from their theoretical sizes, suggesting glycosylation and cleavage [34]. As expected, nucleoprotein was frequently identified and found in bands at 55 kDa, 45 kDa, and 40 kDa. The 55 kDa band in the TMB in-gel staining assay firmly retained hemin. The N protein presence in bands with different predicted molecular masses has been previously noted in SARS-CoV but attributed to protein degradation [35].

Interestingly, N protein was identified in a 46 kDa band of nucleus fraction of infected cells [35]. The transition to a higher molecular mass has been observed for nucleoprotein protein using mass spectrometry [34]. It can be indicative of the complexity of protein-protein binding dynamics in the replication of SARS-CoV-2. Since albumin in culture medium masks low abundance proteins identification [36], we performed a serum depletion followed by affinity chromatography using hemin-agarose beads. This approach increased the overall identification of proteins, and after purification, the N protein was the most abundant protein identified by mass spectrometry in the 45 kDa band. A variety of protein-protein interactions are essential for SARS-CoV-2 replication and viral assembly, such as its structural proteins that have been reported to interact with each other [37]. In the hemin-agarose eluate, the M protein was identified in band 7 at a higher relative molecular mass position than its predicted molecular mass of 25 kDa. It is a transmembrane glycoprotein that comigrated with the other structural proteins Spike and N. While the interaction of the M protein with the N protein has been observed previously and appears to be necessary to viral assembly [38], this comigration was not observed in the total extract of proteins from virally infected Vero cells before the inclusion of the heme-agarose beads. N and S protein are highly immunogenic, unlike M protein [39]. Spike protein appears to bind weakly to hemin beads since no 70 kDa immunoreactive bands were noticed in the eluate fraction.

The heme-binding capacity of viral proteins leads us to question if this could extend to Hb. In the overlay assay, the bands of approximately 150 kDa and 70 kDa, which were exclusively found in the virus-infected cell extract, matched the Spike holoprotein’s predicted size and fragments S1 and S2. The identification of Spike as a significant viral protein in these bands opens Hb’s question and could bind and interact with the RBD region. The Spot-synthesis analysis demonstrated several areas of the RBD of the Spike protein bind Hb, corroborating the molecular docking analysis. The apparent strength of binding according to the strong signals and Hb’s positioning on the Spike protein suggests that its interaction with Spike could hinder viral entry and subsequent replication. In fact, the highest spot signal (448YNYLYRLFRKSNLKP463) was observed within the receptor-binding motif region (RBM; 437-508). This region is responsible for binding to ACE2 and includes hot spot residues that bind Hb (Tyr451 and Leu452). It is essential to highlight that, so far, the most frequent mutations in the RBD region of new variants are located in RBM (Tyr453, Gly476, Phe486, Thr500, and Asn501) but are not present in hot spot amino acids that interact with Hb [40].

Additionally, binding of Spike with hemoglobin may have a role in COVID-19 pathophysiology. Meta-analysis revealed that severe COVID-19 patients had decreased hemoglobin levels, lower RBC count, and higher RDW than moderate COVID-19 cases [41]. Although a combination of events plays a role in COVID-19 hypoxemia [42], the decrease in Hb levels contributes to hypoxia and is related to complications, ultimately leading to multiorgan failure41. Low hemoglobin levels were attributed to hemolytic anemia driven by inflammation and iron dysmetabolism, interfering with erythropoiesis [21,41]; our results demonstrated that viral particles’ binding could also be implicated. In the scenario of hemolytic anemia, an excess of heme/Hb increases ROS levels and tissue damage leading to vascular injury and ferritin overexpression [43]. High ferritin levels are a hallmark of COVID-19, eventually contributing directly to inflammation and lung injury since ferritin is proinflammatory and leads to ferroptosis [7,44]. A recent report found that even after two months from the onset of disease, 30% of patients still presented iron deficiency, hyperferritinemia (38%), and anemia (9.2%) correlated with disease severity [44].

Additionally, free heme and hemoglobin are involved in hemostasis and thrombosis. Hb enhances platelet activation by scavaging nitric oxide23 and also induces platelet aggregation contributing to prothrombotic events [45]. The assembly of viral particles and hemoglobin capping of Spike could contribute to COVID-19 thromboembolic events and aggravate lung injury in critically ill patients. Hemolysis in the intravascular and alveolar spaces results in hemoglobin release that can contribute to organ disfunction [46]; in ARDS, it is proposed that hemoglobin-mediated damage by cell surface receptor binding on the alveolar epithelium is independent of oxidative stress [47].

The ability of porphyrins to interfere in receptor binding was observed through *in vitro* assays. Hemin alone had little effect on viral attachment and replication, suggesting that hemin has little to no direct interactions with the Spike protein or ACE-2. As intracellular concentrations of hemin are highly regulated, and hemin can be exported or degraded by heme-oxygenase11, the absence of an effect is consistent with its physiology. In contrast, exposure of either cell or virus to PpIX dramatically reduced viral load. Similar outcomes for PpIX and verteporfin have been recognized against SARS-CoV-2 since treatment with porphyrins interferes with the ACE2 and Spike that would impair viral entry. Also, these drugs were able to inhibit viral RNA production, suggesting other potential mechanisms of action19. Likewise, our results showed that PpIX at 1µM could inhibit viral replication after 24 h19. PpIX is hydrophobic and could interact with membranes; thus, porphyrin interaction with viral envelope can induce destabilization and oxidation [48].

Pretreatment of viral particles with Hb reduced approximately 50% of both viral replication and adsorption, demonstrating Spike/RBD-ACE-2 fusion impairment. Furthermore, Hb treatment’s effect after viral infection was higher, reducing 99% of viral replication. Modulation of viral replication by extracellular free Hb can occur via prooxidant activity, direct interactions of globin or heme to cell components and signaling pathways, and induction of heme oxygenase [49]. Down-regulating HO-1 is a strategy for the optimization of virus replication and to evade host antiviral mechanisms for hepatitis C virus (HCV), hepatitis B virus (HBV), and Pseudorabies virus (PRV) infection [50–52]. HO-1 is a critical stress-induced enzyme that promotes antioxidant, antiapoptotic, and anti-inflammatory activities via downstream metabolites such as biliverdin and bilirubin [53]. Biliverdin impairs HCV replication by inducing an interferon response [52]. Induction or overexpression of HO-1 inhibits some virus-like influenza [54], HIV [55], human respiratory syncytial virus (RSV) [56], and Zika [57].

Overall, our demonstration that SARS-CoV-2 proteins can bind to heme or Hb may have clinical implications. Mainly, Hb’s interaction with Spike opens new therapeutic perspectives due to significant virus attachment and replication inhibition. Also, this binding could potentially increase or drive hematological disorders and thrombosis observed in severe COVID-19. There are still knowledge gaps on viral-host cell complex interplay and disease pathophysiology, the data presented here will contribute to scientific discussion. More research will be needed to confirm the relevant implications of these heme-protein interactions and correlations with COVID-19 physiopathology.

## Material and methods

### Motifs identification

The MEME Suite server identified the motifs for binding to the heme group on the SARS-CoV-2 nucleoprotein [58]. Protein sequences from 68 crystallized structures with a heme group. Then, 36 Hb sequences deposited in the Uniprot database (www.uniprot.org) were used against 13 nucleoprotein sequences from SARS-CoV-2.

### Molecular docking

#### Receptor-Ligand

Molecular docking assays were employed to predict the binding modes of the Sars-CoV-2 nucleoproteins (PDB code - 6zco; accessed day September 08, 2020 - https://www.rcsb.org/) and ligand heme prosthetic group (Fe-protoporphyrin) complexes by using the DockThor server (https://dockthor.lncc.br/v2/). Structures with positional root mean square deviation (RMSD) ≤ 2 Å were clustered, and results with the most favorable free energy of binding were selected.

#### Protein-Protein

This assay was performed with human hemoglobin (PDB code - 4×0i) and spike protein (PDB code -7kn5), both accessed day September 20, 2020, by using RosettaDock server (http://rosettadock.graylab.jhu.edu). The alpha and beta chains of hemoglobin were assessed in the protein interaction assays and evaluated using the HOTREGION database (http://prism.ccbb.ku.edu.tr/hotregion/index.php).

### Cell culture, virus expansion, and virus tittering

African green monkey kidney (Vero, subtype E6, ATCC^®^CRL-1586^™^) cells were cultured in media consisting of high glucose DMEM complemented with 10% fetal bovine serum (FBS; HyClone, Logan, Utah), 100 U/mL penicillin, and 100 μg/mL streptomycin (Pen/Strep; Thermo Fisher Scientific). Cells were maintained in a humidified atmosphere with 5% CO_2_ at 37 °C. The SARS-CoV-2 used in these studies (GenBank #MT710714) was isolated from a nasopharyngeal swab obtained from a consenting patient with COVID-19, as confirmed by RT-PCR. The virus was expanded in Vero E6 cells at a multiplicity of infection (MOI) of 0.01 according to WHO guidelines that mandate all procedures related to virus cultures be performed in a biosafety level 3 (BSL3) facility.

Virus titer was defined as plaque-forming units (PFU)/mL. Briefly, Vero E6 cells were seeded into 96-well plates at 2 × 10^4^ cells/well for 24 h before exposure to a serial dilution of expanded SARS-CoV-2 for 1 h at 37 °C. A semi-solid high glucose DMEM medium containing 2 % FSB and 2.4 % carboxymethylcellulose was added, and cultures were incubated for 3 days at 37 °C. Then, the cells were fixed with 10 % formalin for 2 h at room temperature. The cell monolayer was stained with 0.04 % solution of crystal violet in 2 0% ethanol for 1h. Virus stocks were stored at -80 °C until use.

### Yield reduction assays and virus titration

Vero cells were seeded into 96-well plates at a density of 2 × 10^4^ cells/well for 24 h at 37 °C before exposure to SARS-CoV-2 at an MOI of 0.01. After a 1 h incubation, the inoculum was removed, and cells were incubated in a medium containing 1 μM of the experimental compounds diluted in DMEM with 2 % FBS. Alternatively, two experimental conditions were performed: i) preincubation of the virus with the compounds (1 μM) for 1 h at 37 °C before their addition to Vero E6 cells (MOI of 0.01) for an additional hour, or ii) preincubation of Vero cells with the compounds (1 μM) for 1 h at 37 °C before their exposure to the virus (MOI 0.01). After 24 h, supernatants were collected for virus titration (PFU/mL) as described above. For virus titration, Vero E6 in 96-well plates (2 × 10^4^ cells/well) were infected with serial dilutions of yield reduction assays’ supernatants containing SARS-CoV-2 for 1 h at 37 °C. A semi-solid high glucose DMEM medium containing 2 % FSB and 2.4 % carboxymethylcellulose was added, and cultures were incubated for 3 days at 37 °C. Then, the cells were fixed with 10% formalin for 2 h at room temperature. The cell monolayer was stained with 0.04 % solution of crystal violet in 20% ethanol for 1 h. The virus titers were determined by plaque-forming units (PFU) per milliliter.

### Adsorption inhibition assays

The virus was incubated with compound (1 µM) for 1 h and then added to monolayers of Vero E6 cells in 48-well plates (5 x 105 cells/well) at an MOI of 0.01 for 1 h at 4 °C. The medium with the virus was removed, and cells washed three times with medium before lysis buffer addition. Total viral RNA was extracted using QIAamp Viral RNA (Qiagen®) according to the manufacturer’s instructions. Quantitative RT-PCR was performed using GoTaq® Probe qPCR and RT-qPCR Systems (Promega) in a StepOne™ Real-Time PCR System (Thermo Fisher). Amplifications were performed as 25 µL reactions containing 1x reaction mix buffer, 50 µM of each primer, 10 µM of the probe, and 5 µL of RNA template. Primers, probes, and cycling conditions followed the recommendations of the Centers for Disease Control and Prevention (CDC) protocol for the detection of the SARS-CoV-2 [59]. A standard curve was included for virus quantification [60].

### SDS-PAGE and hemin-binding blots

Vero E6 cells were infected with SARS-CoV-2 at a multiplicity of infection (MOI) 0.01 for 1 h at 37 °C. The inoculum was removed, and a fresh culture medium was added. Protein extracts (20 µg) were obtained 24 h post-infection by lysing the cell monolayer with lysis buffer (100 mM Tris-HCl pH 8.0, 150 mM NaCl, 10 % glycerol, 0.6 % Triton X100). Protein extracts were electrophoretically separated on a 10 % SDS-PAGE using Laemmli’s buffer [61]. For hemin-binding blots, proteins were transferred to nitrocellulose membrane and then rinsed with Tris-buffered saline [TBS (10 mM Tris-HCl pH 8.0 containing 150 mM NaCl)] plus 0.1% Tween 20 (TBST) followed by 1 h incubation with TBS containing hemin (2 µM). Membranes were subsequently washed three times for 30 min with TBST and revealed with a solution containing 0.1 mg/mL 3,3’ diaminobenzidine (DAB), 0.1 % H2O2, 10 mM HEPES, pH 6.2, and 100 µM CaCl2 overnight at 4 °C in the dark [62]. Alternatively, heme-binding proteins were evaluated by 1 h room temperature incubation of the protein extracts (20 µg) with hemin (300 µM) in 250 mM Tris-HCl, pH 8.0, 5 mM EDTA, and 10% glycerol, followed by SDS-PAGE [63]. Then, gels were washed for 1 h with PBS containing Triton X-100 (2.5 %) and equilibrated for 30 min with sodium acetate 0.5 M (pH 5.0). Heme binding proteins were revealed with 2 mg/mL 3,3’,5,5’-tetramethylbenzidine (TMB) dissolved in 15 mL of methanol and 35 mL of 0.5 M sodium acetate (pH 5.0). Then, 300 μL of 30 % H2O2 solution was added, and the reaction was carried out for 30 min in the dark. After a blue-colored band developing, indicative of a heme-protein complex, the gels were washed with sodium acetate (pH 5.0) and isopropanol (30 %) solution. Protein extract without preincubation with hemin was used as a negative control.

### Hemin and PPIX binding assay

Hemin chloride (10 μM) diluted in NaOH (0.1 M) was added to a quartz cuvette containing 20 μg of protein extract from Vero cells and SARS-CoV-2 infected Vero cells. The absorbance of samples was evaluated at 300–700 nm. Spectroscopic analysis was performed using a SpectraMaxM2e (Molecular Devices). For protoporphyrin IX (PPIX) binding assays, 20 μg of each protein extract was incubated with PPIX solution (1mM in DMSO followed by dilution to 1 µM in phosphate buffer saline pH 7.4) followed by absorbance spectra analysis.

### Protein purification and hemin-agarose binding assay

Protein extract, obtained from infected Vero E6 monolayers 24 h post-infection lysed with 100 mM Tris-HCl pH 8.0, 150 mM NaCl, 10 % glycerol, 0.6 % Triton X-100, was initially subjected to affinity chromatography using a protein-A/anti-BSA mAB matrix (Sigma-Aldrich) to remove excess albumin (a heme-binding protein contaminant). Then, hemin-agarose was used to isolate heme-binding proteins. Briefly, 200 mL of hemin-agarose (Sigma-Aldrich) was washed three times in 1 mL of 100 mM NaCl, 25 mM Tris-HCl (pH 7.4) 5 min, and centrifugated at 700 *g*. Next, hemin-agarose was incubated for 1 h at 37 °C, under agitation, with SARS-CoV-2 protein extracts (800 µg). The unbound proteins were removed by washing three times with equilibration buffer, and beads incubated for 2 min with elution solution (2 %, SDS, 1 % β-mercaptoethanol in 500 mM Tris HCl, pH 6.8) followed by boiling at 100 °C for 5 min [64]. Total extract, unbound (supernatant) fraction, washing fraction, and hemin-agarose bound proteins were separated by 10 % SDS–PAGE gels and stained with Coomassie Blue R-250 or transferred to a nitrocellulose membrane (subsequently blocked with TBS-T (Tris-buffer saline, 0.1% Tween 20, pH 7.5) and 5% defatted milk. Immunostaining was later performed by incubating membranes overnight at 4 °C with a pool of COVID-19 convalescent serum (1:200). Then, membranes were washed and incubated with HRP-conjugated anti-human IgG antibody (Sigma-Aldrich; 1:10.000) followed by chemiluminescence detection.

### Overlay assay

Protein extracts (20 µg), obtained as described above, were separated by 10 % SDS– PAGE and transferred to a nitrocellulose membrane (Bio-Rad). Haptoglobin (Sigma-Aldrich) was used as a control. The membrane was blocked with 2 % BSA in TBS-T buffer, then incubated overnight at 4 °C after adding 10 µg/mL of human hemoglobin (Hb). After washing with TBS-T, the membrane was incubated with an anti-Hb antibody (1:5.000) (Sigma-Aldrich). The antigen-antibody complex was revealed by chemiluminescence.

### Spot-Synthesis

The DNA sequence of the Spike protein (P0DTC2) receptor-binding domain was retrieved from Uniprot. A library of 15 amino acid peptides with a 5-amino acid overlap was designed to represent the entire coding region (319-541 aa) of RBD and automatically synthesized onto cellulose membranes using an Auto-Spot Robot ASP222 (Intavis, Koeln, Germany) according to the SPOT synthesis protocol [65,66]. Briefly, membranes containing the synthetic peptides were washed with TBST (and then blocked with TBS-T containing 1.5 % BSA under agitation for 2 h at room temperature. After extensive washing with TBS-T (Tris-buffer saline, 0.1 % Tween 20, pH 7.0), membranes were incubated overnight with Hb (5 µg/mL) dissolved in TBST + BSA (0.75 %). After incubation, membranes were washed with TBS-T, followed by additional incubation with anti-human Hb antibody for 90 min. Subsequently, the membrane was washed with TBS-T and incubated for 90 min with anti-rabbit IgG antibody conjugated to alkaline phosphatase (Sigma Alrich), diluted 1:5.000 in TBS-T solution containing 0.75 % BSA. Washes were performed with TBS-T followed by the addition of substrate for chemiluminescent alkaline phosphatase Tropix® was added. Next, membranes were washed three times with TBS-T, and then the buffer was exchanged to CBS (50 mM citrate-buffer saline) before the addition of the chemiluminescent enhancer Nitro-Block II. The chemiluminescent substrate Super Signal R West Pico was applied, and signals were immediately detected. A digital image file was generated, and the signal intensities quantified using TotalLab (Nonlinear Dynamics, USA) software^66^. The spots with signal intensity greater than or equal to 50% of the highest signal value obtained in all membrane spots were considered to identify possible binding motifs.

### In-Gel Trypsin Digestion of Proteins

Protein spots were excised from gels using sterile stainless steel scalpels, transferred to 0.5 mL tubes, and cut into smaller pieces. In-gel digestion with trypsin (Promega V511A) was performed according to the literature [67] with modifications described elsewhere [68]. Protein reduction was performed by the addition of 100 μL of 65 mM DTT for 30 min at room temperature, followed by alkylation with 100 μL of a 200 mM iodoacetamide solution for 30 min, in the dark, at room temperature. After washes and trypsinization, the final 80 μL peptide-containing samples were concentrated by vacuum centrifugation to approximately 20 μL and stored at −20°C until mass spectrometric analysis. Gel pieces from a “blank” region and the BSA molecular mass marker were negative and positive controls, respectively.

### Identification of proteins by mass spectrometry

The tryptic digests were analyzed in three technical replicates by reversed-phase nanochromatography coupled to high-resolution nanoelectrospray ionization mass spectrometry. Chromatography was performed using a Dionex Ultimate 3000 RSLCnano system coupled to the HF-X Orbitrap mass spectrometer (Thermo Fischer Scientific). Samples (4 µL per run) were initially applied to a 2 cm guard column, followed by fractionation on a 25.5 cm PicoFritTM Self-Pack column (New Objective) packed with 1.9 μm silica, ReproSil-Pur 120 Å C18-AQ (Dr.Maisch/ Germany). Samples were loaded in 0.1 % (v/v) formic acid (FA) and 2 % acetonitrile (ACN) onto the trap column at 2 μL/min, while chromatographic separation occurred at 200 nL/min. Mobile phase A consisted of 0.1% (v/v) FA in water, while mobile phase B consisted of 0.1% (v/v) FA in ACN. Peptides were eluted with a linear gradient from 2 to 40% eluent B over 32 min, followed by up to 80% B in 4 min. Lens voltage was set to 60 V. Full scan MS mode was acquired with a resolution of 60,000 (FWHM for *m/z* 200 and AGC set to 3×10^6^). Up to 20 most abundant precursor ions from each scan (*m/z* 350-1,400) were sequentially subjected to fragmentation by HCD. Fragment ions were analyzed at a resolution of 15,000 using an AGC set to 1×10^5^. Data were acquired using Xcalibur software (version 3.0.63).

### Peptide identification and protein inference

All MS/MS spectra were analyzed using PEAKS Studio X Plus (Bioinformatics Solutions, Canada). Peptide identification was performed against *Chlorocebus sabaeus* reference proteome at the UNIPROT database under ID UP000029965, plus the SARS-CoV-2 reference proteome at the same database UP000464024 (downloaded July 03, 2020). Data refinement applied the precursor correction (mass only). Next, PEAKS *de novo* analysis was run assuming trypsin digestion, with a fragment ion mass tolerance of 0.02 Da and a parent ion tolerance of 10 ppm. Cysteine carbamidomethylation (+57.02 Da) was set as fixed modification; a maximum of 2 variable shifts per peptide was allowed. PEAKS DB analysis was performed using these same parameters, plus the possibility of up to two missed enzyme cleavages and nonspecific cleavage at both sides of the peptides. Finally, post-translational and other possible modifications were searched using the PEAKS PTM algorithm, with the same parameters described above, against a protein subdatabase composed only by protein entries found by the previous PEAKS De Novo and PEAKS DB searches. False discovery rates (FDR) were estimated through the PEAKS decoy fusion approach. A peptide-spectrum match FDR of 0.1 % and protein identifications with at least 2 unique peptides were the criteria used to establish FDR values at peptide and protein levels smaller than 1 %.

### Data analysis

Graphs were prepared, and statistics were performed using R software 4.0.0 and GraphPad Prism version 9.

## Author Contributions

GCL, FSS, SGS, and TMLS conceived and designed the proposal. GCL, MCSP, RHV, TMLS, SGS, and DWP wrote the paper. GCL, FSS, CQS, SSGD, NFR, JRT, and PNP performed *in vitro* experiments and processed the data. FSS, CRA, and NC performed *in silico* and docking analysis. MROT and RHV performed mass spectrometry and data analysis. Contributed with reagents/materials/analysis: SGS, TMLS, MCSP, MROT, and RHV. All authors have read and agreed to the published version of the manuscript.

## Funnding

SGS received support from FIOCRUZ (INOVA VPPCB-007FIO-18-2-21 and VPPIS-005FIO-20-2-51), Carlos Chagas Filho Foundation for Research Support of the State of Rio de Janeiro/FAPERJ #(grant E-26 110.198-13), and the Brazilian Council for Scientific Research/CNPq #(grant 467.488.2014-2 and #301744/2019-0).

DWP received support from FIOCRUZ (INOVA VPPCB-005-FIO-20).)

TMLS received support from FIOCRUZ (INOVA B3-Bovespa)

## Acknowledgments

SGS, CRA, MCSP and RHV are Post Doctorate fellows from CNPq.

GCL is Post Doctorate fellow from CINCT-IDPN/FAPERJ (E-26/201.848/2020)

FSS is Post Doctorate fellows from CAPES.

CQS, NFR and JTR are postdoctoral fellows from CAPES/CDTS and Inova Program.

SSGD is a PhD student from Oswaldo Cruz Institute/Fiocruz.

---

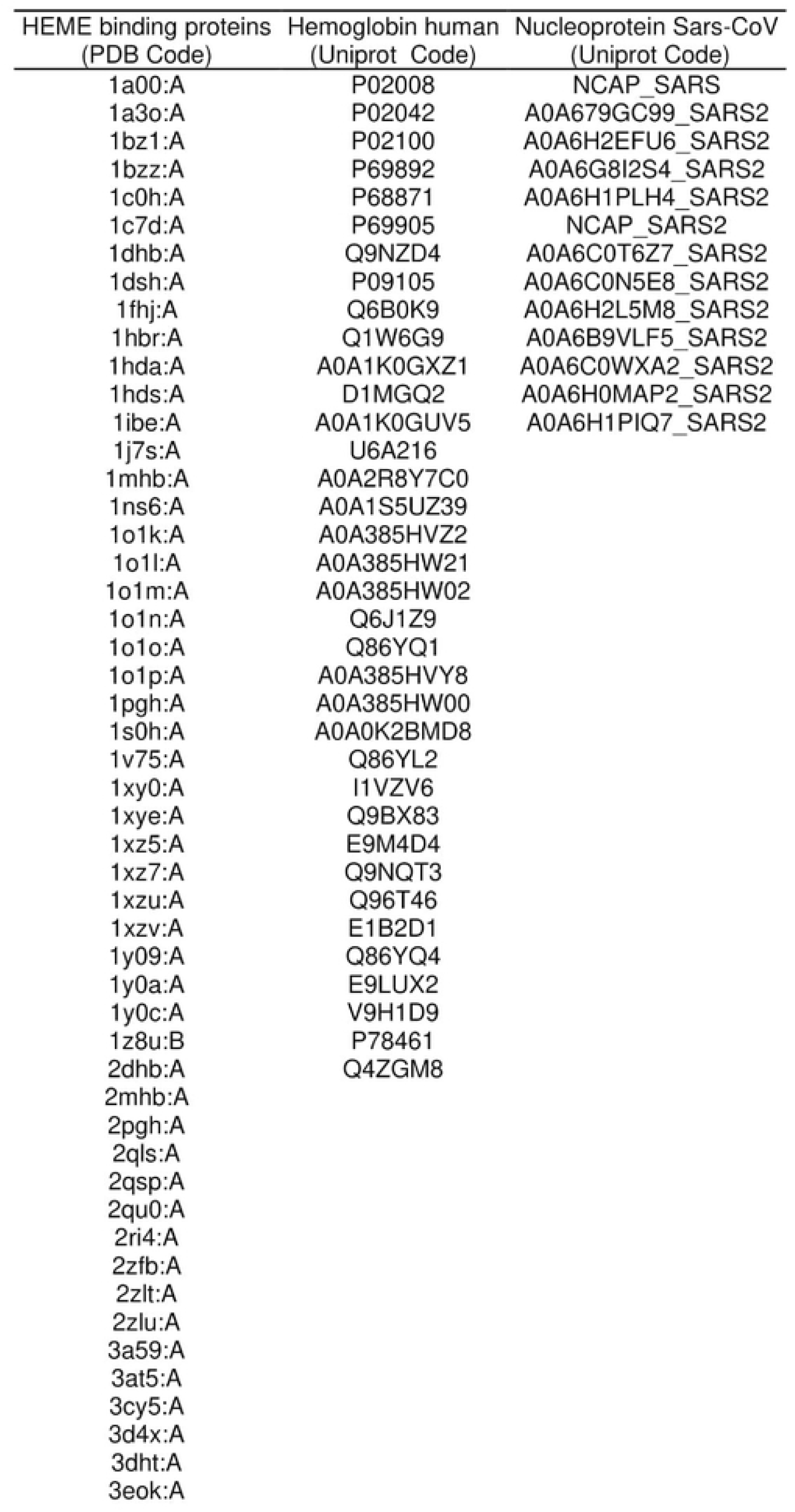
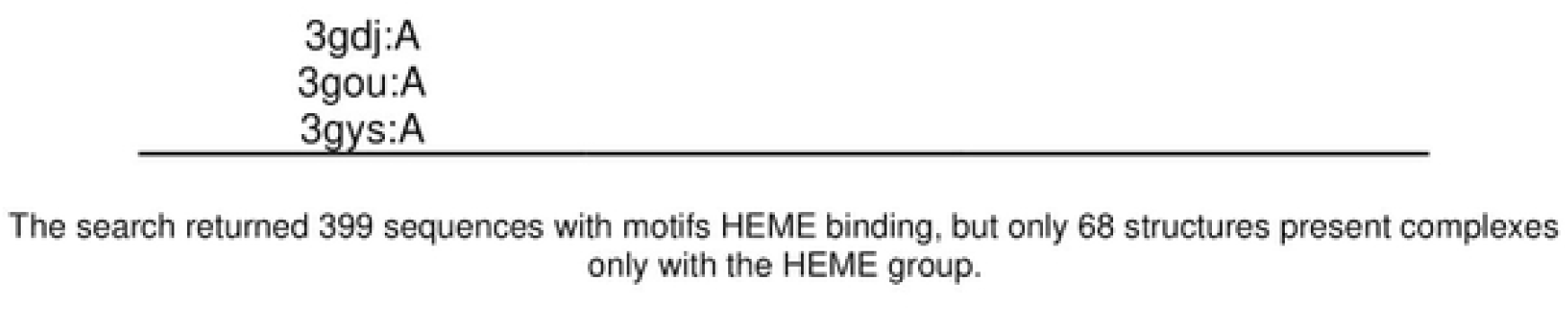

| Protein | Residue | Complex<br>ASA<sup>b</sup> | Monomer<br>ASA<sup>b</sup> | Pair<br>Potential | Hotspot<br>Status <sup>a</sup> | Hotregion<br>Status | Complex<br>ASA<sup>b</sup> | Monomer<br>ASA<sup>b</sup> |
| --- | --- | --- | --- | --- | --- | --- | --- | --- |
| HB_A | LEU35 | 26.04 | 73.04 | 17.36 | NH |  | 46.51 | 130.4 |
| HB_A | PRO38 | 42.98 | 61.77 | 2.43 | NH |  | 58.51 | 84.09 |
| HB_A | LYS41 | 0 | 0 | 13.92 | NH |  | 0 | 0 |
| HB_A | THR42 | 0 | 0 | 0 | NH |  | 0 | 0 |
| HB_A | PRO45 | 0 | 0 | 4.99 | NH |  | 0 | 0 |
| HB_A | HIS46 | 0 | 0 | 9.17 | NH |  | 0 | 0 |
| HB_A | PHE47 | 0 | 0 | 14.04 | NH |  | 0 | 0 |
| HB_A | ASP48 | 0 | 0 | 19.97 | H | 2 | 0 | 0 |
| HB_A | SER50 | 0 | 0 | 17.74 | NH |  | 0 | 0 |
| HB_A | HIS51 | 0 | 0 | 19.4 | H | - | 0 | 0 |
| HB_A | GLY52 | 0 | 0 | 11.09 | NH |  | 0 | 0 |
| HB_A | SER53 | 0 | 0 | 7.25 | NH |  | 0 | 0 |
| HB_A | ALA54 | 0 | 0 | 7.56 | NH |  | 0 | 0 |
| HB_A | GLN55 | 0 | 0 | 12.46 | NH |  | 0 | 0 |
| RBD_C | GLU340 | 47.25 | 67.63 | 7.68 | NH |  | 81.38 | 116.5 |
| RBD_C | ALA344 | 0 | 0 | 7.56 | NH |  | 0 | 0 |
| RBD_C | THR345 | 0 | 0 | 6.17 | NH |  | 0 | 0 |
| RBD_C | ARG346 | 0 | 0 | 27.67 | H | 2 | 0 | 0 |
| RBD_C | PHE347 | 0 | 0 | 22.7 | H | 2 | 0 | 0 |
| RBD_C | ALA348 | 0 | 0 | 17.73 | NH |  | 0 | 0 |
| RBD_C | SER349 | 0 | 0 | 12.03 | NH |  | 0 | 0 |
| RBD_C | TYR351 | 0 | 0 | 24.76 | H | - | 0 | 0 |
| RBD_C | ALA352 | 0 | 0 | 16.19 | NH |  | 0 | 0 |
| RBD_C | TRP353 | 0 | 0 | 16.59 | NH |  | 0 | 0 |
| RBD_C | ASN354 | 0 | 0 | 10.77 | NH |  | 0 | 0 |
| RBD_C | ARG355 | 0 | 0 | 15.53 | NH |  | 0 | 0 |
| RBD_C | ASN450 | 0 | 0 | 13.12 | NH |  | 0 | 0 |
| RBD_C | TYR451 | 0 | 0 | 31.74 | H | 1 | 0 | 0 |
| RBD_C | LEU452 | 0 | 0 | 33.83 | H | 1 | 0 | 0 |
| RBD_C | ARG466 | 26.46 | 58.18 | 12.34 | NH |  | 63.18 | 138.9 |
| RBD_C | ILE468 | 22.33 | 56.43 | 10.14 | NH |  | 39.11 | 98.82 |
| RBD_C | LEU492 | 0 | 0 | 22.69 | H | 1 | 0 | 0 |

<sup>a</sup> NH denotes non hot spot and H denotes hot spot and <sup>b</sup> Solvent Accessible Surface Area (ASA)

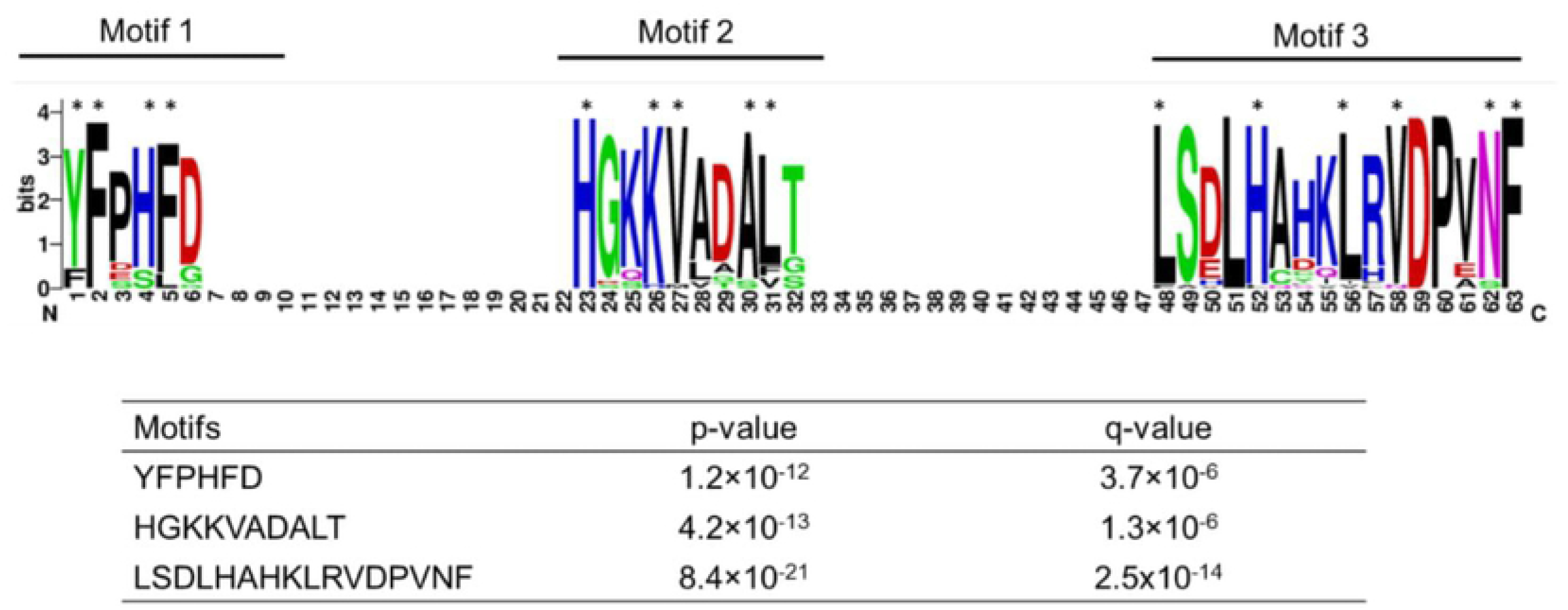

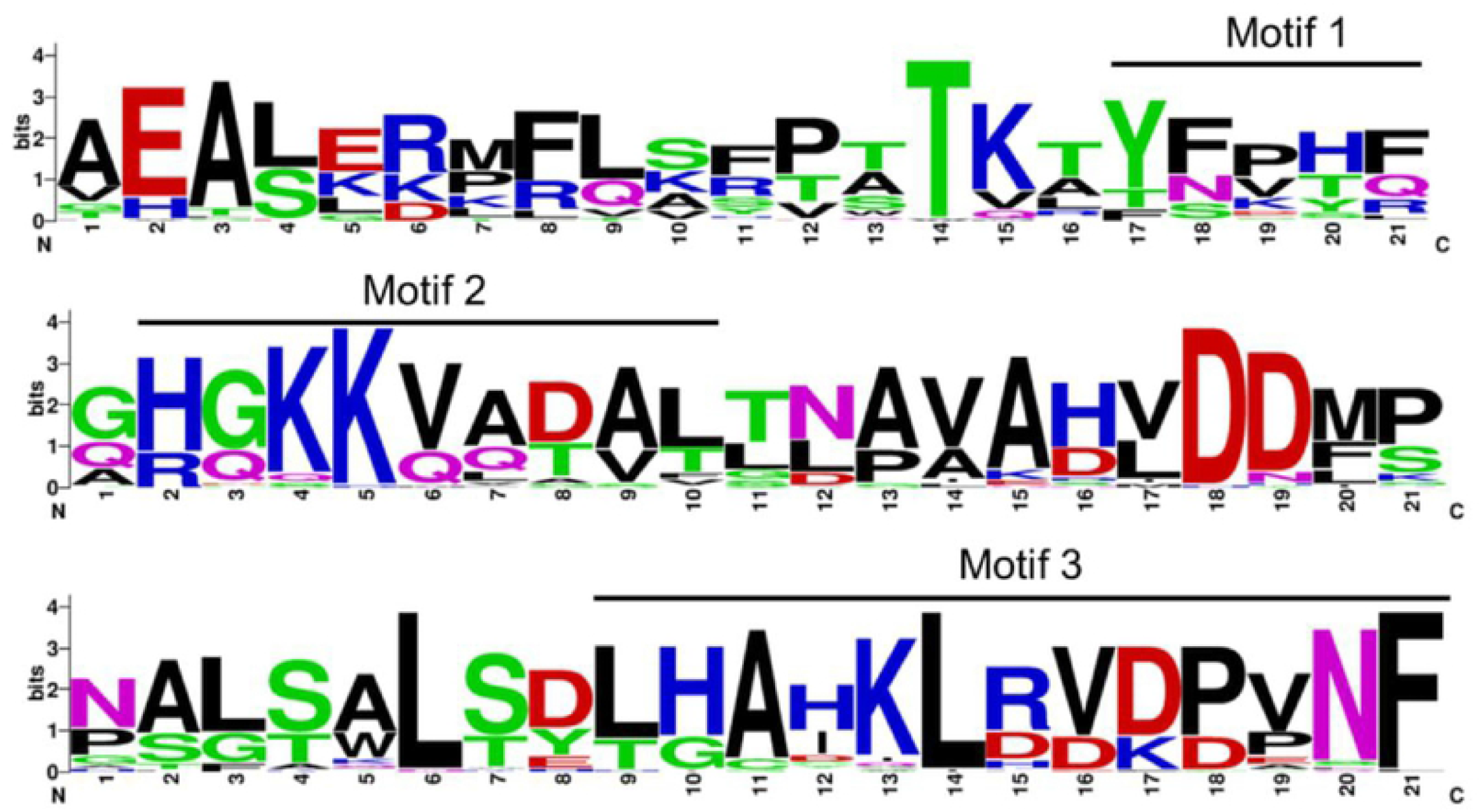

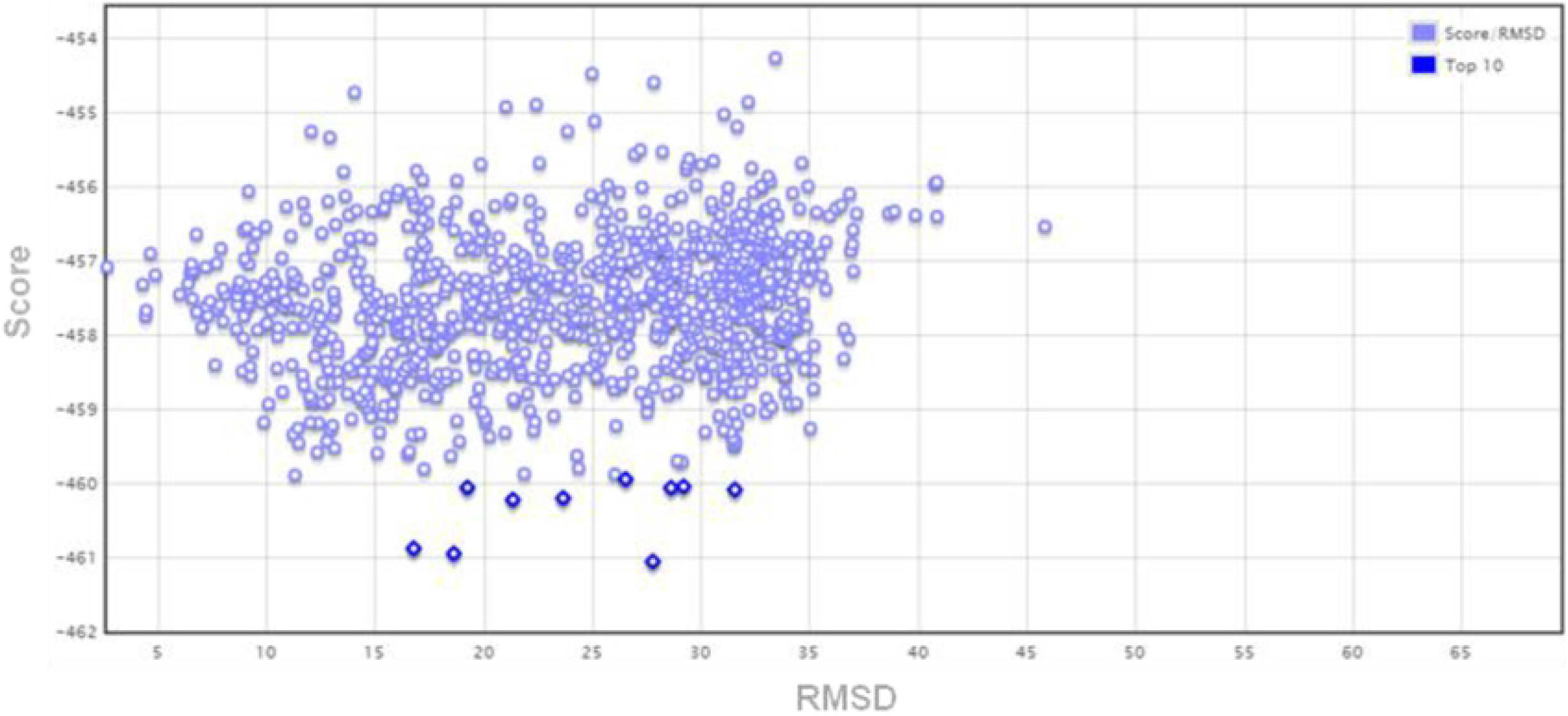

